# Mechanical competition alters the cellular interpretation of an endogenous genetic programme

**DOI:** 10.1101/2020.10.15.333963

**Authors:** Sourabh Bhide, Denisa Gombalova, Gregor Mönke, Johannes Stegmaier, Valentyna Zinchenko, Anna Kreshuk, Julio M Belmonte, Maria Leptin

## Abstract

The intrinsic genetic programme of a cell is not sufficient to explain all of the cell’s activities. External mechanical stimuli are increasingly recognized as determinants of cell behaviour. In the epithelial folding event that constitutes the beginning of gastrulation in *Drosophila*, the genetic programme of the future mesoderm leads to the establishment of a contractile actomyosin network that triggers apical constriction of cells, and thereby, tissue folding. However, some cells do not constrict but instead stretch, even though they share the same genetic programme as their constricting neighbours. We show here that tissue-wide interactions force these cells to expand even when an otherwise sufficient amount of apical, active actomyosin is present. Models based on contractile forces and linear stress-strain responses do not reproduce experimental observations, but simulations in which cells behave as ductile materials with non-linear mechanical properties do. Our models show that this behaviour is a general emergent property of actomyosin networks [in a supracellular context, in accordance with our experimental observations of actin reorganisation within stretching cells.

## Introduction

Epithelial tissues are shaped during animal development by changes in the geometry, number or relative positions of their constituent cells. Cells change their shape by actively generating intracellular forces or by passively responding to external forces, from within the organism, such as neighbouring cells, or by forces from outside the body^1–4^.The actomyosin meshwork underlying the plasma membrane is the major source of morphogenetic forces^5–7^ which can be transmitted over larger, supracellular distances via cell junctions. The functioning of the cytoskeleton itself can be influenced by external mechanical forces^8^. In some systems, we are beginning to understand how forces act on a tissue scale^9^, but we know much less about the interplay of active forces and passive deformation and their genetic and molecular basis. Understanding the actomyosin contraction patterns in the individual cells that make up a tissue is unlikely to be sufficient to explain all the force changes and deformations within the entire tissue.

One example of epithelial morphogenesis is the formation of the ventral furrow during *Drosophila* gastrulation, an epithelial folding event that internalizes the future mesoderm, driven by active forces generated in an autonomous manner in the central part of the mesoderm^10^. Many studies have focused on these cells and their contractile actomyosin meshwork. We understand the major mechanisms that act within each cell: the proteins that are specifically activated in these cells change the location of the adherens junctions and recruit an active actomyosin meshwork to the apical cell cortex, which undergoes a series of pulsatile contractions until the apical surface is fully constricted^11–14^.

To allow the furrow to internalize the mesoderm without causing disruptions elsewhere in the embryo, other parts of the embryonic epithelium obviously must respond or contribute to the movement. The cells outside the mesoderm appear not to contribute actively to furrow formation^15^, but their compliance is later required for the furrow to invaginate fully^16^. The most important cells that enable the furrow to form are the mesodermal cells adjacent to the initial indentation. While central cells constrict, lateral cells expand their apical surfaces^16,17^.

In spite of their distinct behaviours, the constricting and expanding cells of the mesoderm share the same developmental program. They express the same genes, albeit with quantitative differences, but no known genes are absolutely restricted to one or other population^18,19^. There is a graded expression of important gene products from the centre to the edges of the mesoderm (Fig. 1A), in particular for the genes necessary for myosin activation (*fog, t48* and *mist*) and junction remodelling (*traf4*) which are deployed under the control of the dorsal-ventral patterning system^20–27^. While their quantitative differences have prompted the question whether the two populations should be considered distinct ‘subdomains’, each relevant gene has a different expression boundary, so they together cannot be seen as defining a genetic domain^27^.

**Fig. 1.**
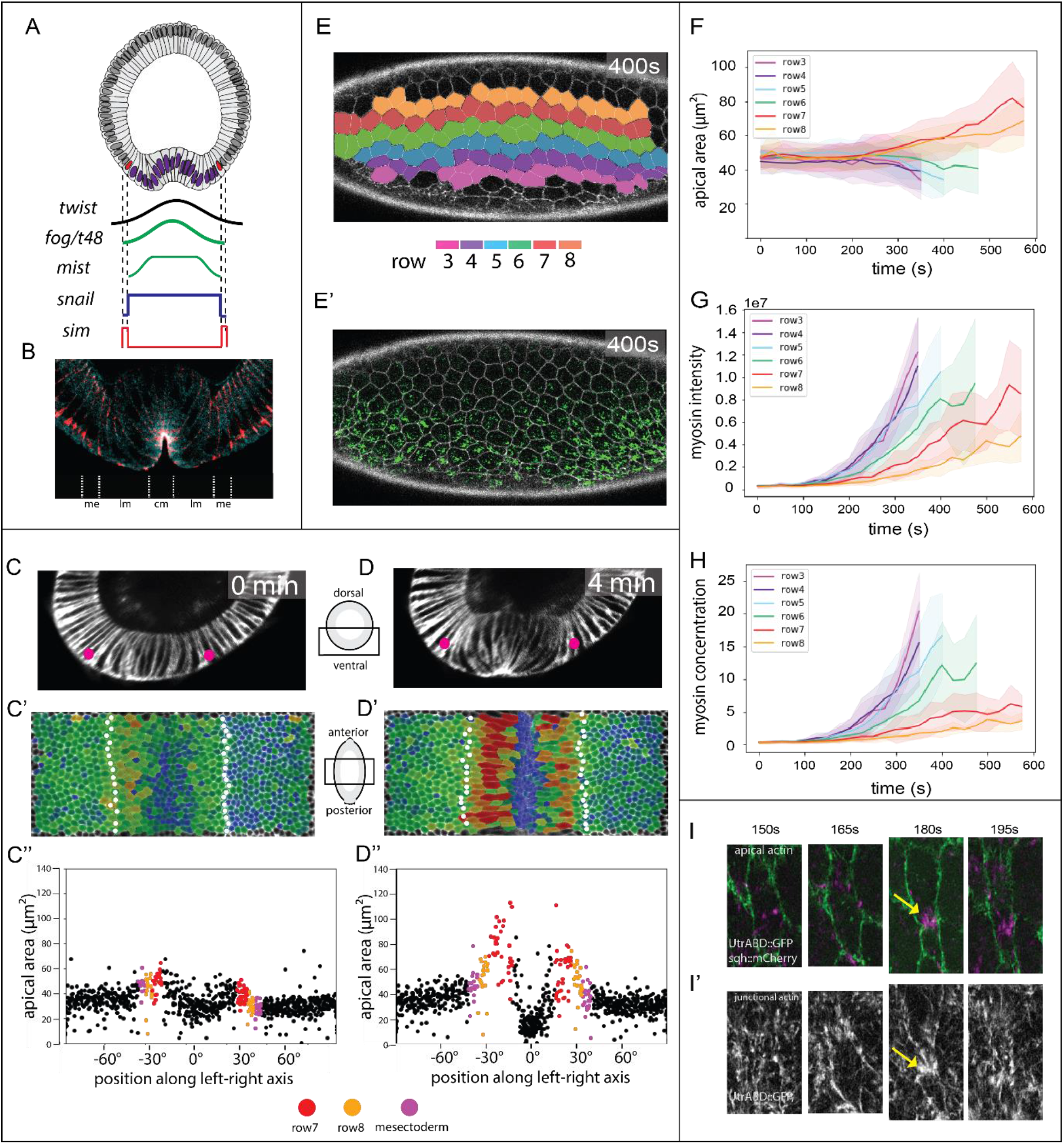
Cell activities during ventral furrow formation. (A) Genes expressed ventrally at the onset of gastrulation. Top: Diagram of a cross-section through an embryo at the beginning of gastrulation. Mesodermal nuclei expressing Snail: blue, mesectodermal nuclei with single-minded: red. Bottom: Schematic of gene expression levels. Twist (black) and Snail (blue) regulate the genes that control shape changes (*fog, T48, mist*). (B) Section of an embryo stained for beta-catenin/armadillo to visualize adherens junctions (pink) and myosin (blue). Junctions in the central (cm) and lateral (lm) mesoderm are apical, the mesectodermal (me) cell has one on apical and one subapical junction, ectodermal junctions are subapical^25^. (C, −D) Cross-sectional views at two time points from a MuVi-SPIM recording of an embryo expressing GAP43::mCardinal (membrane). Pink dots: mesectoderm. timeseries in Suppl. Fig.1 (C’, D’) Apical surface ‘peels’ with colour-coded apical cell areas. Mesectoderm: white dots. (C”, D”) Apical area from C’ and D’ plotted against cell position (0° is the ventral midline). Each dot represents one cell. Colour-code for rows 7, 8 as in E, mesectoderm magenta. (E) Ventro-lateral views of a confocal recording of an embryo expressing Spider::GFP (white) and sqh::mCherry (green) at a confocal Z-plane 3μm below the surface (Suppl. Movie 2). (E’) Cells were segmented using Spider::GFP and assigned to colour-coded rows. (F-H) Apical areas, total myosin intensity and myosin concentration plotted per row against time (mean and standard deviation). Tracks for ventral rows stop early because the cells are lost from the imaging plane. (I-I’) Example of a lateral mesodermal cell at four time-points in an embryo expressing utrABD::GFP (subapical for cell outlines in I, green; apical in I’; white) and sqh::mCherry (magenta) during formation of a myosin focus. Arrow: local cortical deformation.

Current models for cell shape determination in the ventral furrow^28–32^ assume that changes in apical surface area correlate with the force generated by contractile actomyosin. In the first phase of invagination, the degree of apical constriction mirrors the graded distribution of apical myosin, with absence of myosin having been correlated with lack of constriction of lateral cells^16,33,34^.

It is not clear by what mechanism the quantitative differences in gene expression can cause dramatic qualitative differences in cell behaviour: any two immediately adjacent cells in the mesoderm primordium have similar gene expression profiles. Thus, in the absence of any known genetic correlations for the pronounced differences, there must be other explanations for how these behaviours arise. Specifically, we need explanations for how the smooth and graded differences in expression levels of effector molecules is converted into a step difference in cell behaviour.

We compare here in a quantitative manner the cellular activities in the mesoderm, contrast them with existing models, and propose and test a new model that explains qualitative differences in cell behaviour. Our results suggest that two distinct cell behaviours emerge not from strict differences in genetic control, but from tissue-wide mechanical interactions.

## Results

### Cell shape evolution across mesoderm and neighbouring populations

Analyses of shape changes in the prospective mesoderm (hereafter simply called ‘mesoderm’) often focus on the 10-cell-wide central band of cells that form the initial furrow. The lateral cells are less well studied, partly because the forces for folding are generated in central cells, but also because their rapid displacement and extreme shape changes make them difficult to image^35^. We extracted faithful two-dimensional views of the apical surface of the entire mesoderm (surface ‘peels’^36^ Suppl. Fig. 1A-C) for quantitative analysis. The breadth of the mesoderm varies along the AP axis and between embryos; we therefore define the cell rows operationally from row 1 at the midline to row 8 as the outer row adjacent to the mesectoderm; Suppl. Fig. 1).

Furrow formation starts with cells in rows 1 - 6 constricting in a stepwise and stochastic manner^12,37^. The last cells to constrict are those in row 6, while rows 7 and 8 expand their surfaces anisotropically^21–23,33,35^, stretching towards the midline. Mesectodermal cells also stretch slightly, but beyond them the ectoderm remains inert. Thus, mesodermal cells can either constrict or stretch, with initially indistinguishable neighbours in rows 6 and 7 taking on dramatically different developmental paths. In addition, rows 7 and 8 do not respond equally to the force from the centre. Row 7 expands first and most strongly, followed by row 8 and finally the mesectoderm (Fig. 1C-D).

Theoretical models and simulations based on bell-shaped contractility gradients create epithelial shape changes with highly constricted cells in the centre and cell sizes increasing in a graded manner with distance from the centre^16,31–33,38^. Inverted patterns of stretching have so far been obtained in computational models only for cells without contractility^31,39,40^. To investigate this inconsistency, we examined actomyosin in lateral cells.

### Actomyosin gradient as a predictor for cell shape behaviour

F-actin is present in two distinct but interacting pools with different morphological functions in the early embryonic epithelium: a fine meshwork underlying the apical cortex, and a large pool associated with apical junctions and baso-lateral cell membranes^12,31,41–45^. Junctional actin is reduced in the mesoderm in unison with the relocation of adherens junctions before shape changes begin^44,46,47^ (Suppl. Fig. 2). The apical meshwork changes along the entire dorso-ventral axis around the time of gastrulation but remains present during furrow formation as a fibrous network both in central and lateral mesodermal cells^16^ (Suppl. Fig. 2I-K).

We focused our further analyses on myosin, on which the contractile forces in the mesoderm depend. The amount of myosin regulatory chain (encoded by the gene *sqh* in *Drosophila*) within the apical cortex has been used as a proxy for the contractile actomyosin meshwork^12,48–51^. When the central cells begin to constrict, practically no apical myosin is seen in the lateral cells^33^ (Fig. 1D-E). Levels rise over the next few minutes, reaching values seen in central cells at earlier points, when the cells constrict. For example, the level in row 7 at 525sec resembles that in rows 3 and 4 at 325 sec. We also calculated the concentrations, and still find that row 7 at 525 sec reaches similar concentrations as rows 3 to 5 at 275 sec (Fig. 2F). Thus, apical myosin levels alone are not sufficient to explain why lateral cells do not constrict.

**Fig. 2.**
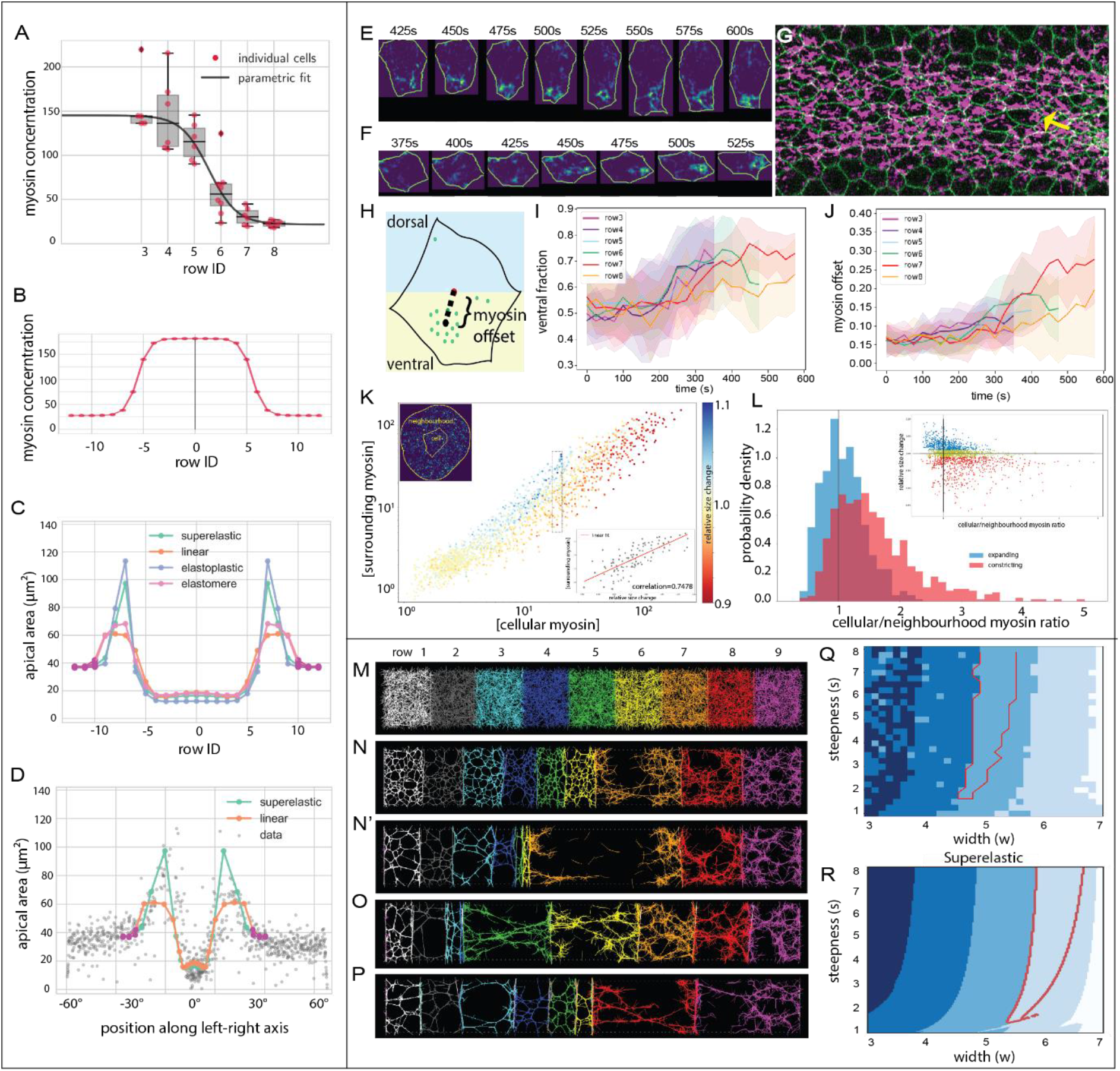
Computational models and myosin distribution. (A-D) Viscoelastic model of a line of cells. (A) Polynomial fit to the myosin concentration per row, measurements from embryo 1. (B) The model is driven by an explicit contractility value for each cell. (C) Final cell lengths for linear elastic, elastomeric and elastoplastic spring constants. Magenta dots represent the stiffer ectoderm. (D) Values for two curves from (C) superimposed on measured cell sizes. The point for each value is shifted along the x-axis from the starting point represented in (C) to the position where each cell row has moved at this time point. (E) Example of myosin dynamics in a single cell from row 8. Cell contours and myosin signal pixels were isolated using individual cell segmentation masks. Myosin intensity values increase from blue to yellow. (F) Example of an incompletely constricted central mesodermal cell (arrow in G). (G) Embryo 2, ventral view. Green: membranes; magenta: myosin. (H) Representation of myosin spatial distribution in a cell. ‘Offset’ is the Euclidean distance between the cell centroid and the intensity-weighted centroid of the myosin signal; ‘DV asymmetry’ is the ratio of myosin pixels in the ventral half of the cell to the total number of myosin pixels in the cell. (I) Average proportion of myosin in the ventral half of the cell, plotted over time for each row. (J) Offset of myosin centroid from cell centroid, average per row. (K) Myosin concentration within a cell plotted versus surrounding myosin concentration in a ring around the cell (radius of 70 pixels (~8.5 μm) from each point of the cell periphery; top left inset), with the change in cell size over two consecutive frames indicated in colour. All segmented cells at 25 time points from movie 5 are represented. The bottom right inset shows the cells with internal concentrations at values between 18 and 22 (boxed in the main plot) with surrounding concentration plotted against size change. (L) Change in cell size compared to the ratio of cell-intrinsic over surrounding myosin concentration. The top right insert shows a plot of all individual cells, with colour illustrating the bins used for the main density histogram (blue: expanding; red: contracting; yellow: no significant change, not represented in the histogram). All cells with concentrations above 45 constrict, regardless of the levels in surrounding cells. The proportion of expanding cells is greater at low intrinsic-to-surrounding levels, and is highest when this ratio drops below one (i.e. surrounding cells have more myosin). (M-Q) Microscopic model of a line of cells with a contractile actomyosin meshwork. (M) Initial condition of the system with randomly distributed actin, crosslinkers and myosin motors within each cell (shown with different colors). (N-N’) Example of a simulation with myosin profile that qualitatively reproduces experimental results. (O-P) Examples of simulations where the myosin profile was wider (M) or shorter (N). (Q-R) Parameter map for myosin concentration curves with varying peak widths and steepnesses for microscopic (Q) and visco-elastic (R) with super-elastic models. Blue shades: number of expanding cells. Red outline: conditions where the three right cells expand with an inverted pattern of stretching that qualitatively matches experiments.

Another possibility is that in spite of having sufficient myosin, lateral cells cannot assemble a functional contractile meshwork. Epithelial apical actomyosin meshworks normally show a strong dynamic behaviour characterised by fluctuations or ‘pulses’ of myosin foci that correlate with periods of apical constriction^12,48,52^. We see myosin foci forming, moving and disappearing in lateral cells in a similar manner as in central cells (Fig. 2G). Myosin pulses in lateral cells have been characterised as less persistent^47^, but they are nevertheless able to pull on nearby plasma membrane, thereby narrowing the cell (Suppl. Fig. 3), indicating an active, force-generating actomyosin meshwork. Thus, in this regard lateral cells are not qualitatively different from central cells.

### Visco-elastic model of the mesoderm

Taking into account the myosin levels in lateral cells, we explored in a computational model whether a simple contractility gradient could explain the bifurcation into constriction and expansion, the inverted pattern of stretching and the apical size ratios. With a mathematical description of our myosin measurements per cell row, we modelled the mesoderm and mesectoderm as a line of 19 visco-elastic “cells” with a given stress-strain response, bordered by three stiffer ‘ectodermal’ cells on each side. Each “cell” changes size based on the forces acting on its boundaries, which in turn depend on the difference of the myosin levels in the cells on either side of the boundary (Suppl Fig. 4). The simulation showed constriction in central cells and stretching in lateral cells, but not with the pattern of size ratios observed in the embryo. This might be explained by inaccuracies in our myosin measurements, but systematically varying the width and steepness of the myosin profile also did not yield outputs corresponding to the *in vivo* data, nor did changes in the slope of the stress-strain curves.

We therefore tested whether the assumption of a linear stress-strain response in the cells was wrong, as also seems to be the case in other instances^8, 53, 66, 67^. We considered four classes of non-linear stress-strain responses: superelastic (like nickel-titanium alloys) with strain-softening beyond the proportional limit followed by strain-hardening while remaining elastic; elastoplastic (like aluminium), with a similar stress-strain relationship but permanent deformation (yielding); elastomeric (like rubber or silicon), with a decrease in stiffness after the proportional limit, but no strain-softening; and a stiffening model (like biopolymer networks), with increased stiffness after the proportional limit (Suppl Fig. 5B). Unlike the linear models, non-linear models with strain-softening (superelastic- and elastoplastic) reproduced the stretching pattern of lateral cells for a wide range of myosin profiles (Fig. 2, Suppl Fig. 5). While inert materials and cultured cells can respond to strain by stiffening ^64–67^, simulations with strain-stiffening curves did not reproduce our *in vivo* observations. These results led us to re-examine the actomyosin meshwork in lateral cells since a strain-softening would most likely manifest as permanent or reversible reorganisations of the cytoskeletal network.

### Organisation of actomyosin networks in lateral cells

We had noticed that local constrictions in lateral cells occurred primarily in the AP axis (Suppl Fig. 5), pointing to a possible role for overall actomyosin distribution. We analysed the distribution of apical myosin and found preferential segregation towards the ventral side in each cell (Fig. 2F). While the asymmetry is visible in all cell rows, there are larger areas without myosin and the distance of displacement is greater in lateral cells (Fig. 2G-J). This uneven distribution may reflect the strain-softening or yielding behaviour predicted necessary by the model. This resembles the asymmetric distribution of Rho in expanded central cells in *concertina* mutants, which has been proposed as an explanation for the cells’ inability to overcome the expansile forces acting on them and constrict^54^. The reason for the asymmetry may be the myosin gradient. For every cell along the gradient the ventral neighbour constricts earlier than its dorsal neighbour. Recent simulations showed that the ability of the cell cortex to yield to contractile forces feeds back on the orientation of the contractile network, which becomes depleted near ‘softer’ and enriched near ‘stiffer’ membranes^55^. In mesodermal cells, the least yielding should be the ventral side, which experiences stronger forces from the ventral neighbour than the other side does from the dorsal neighbour. If the extent to which a cell at any moment expands or shrinks is influenced by its neighbours, then the differential concentration of myosin within the cell and its surrounding should correlate with the cell’s size changes. We therefore compared these parameters and did indeed find such a correlation (~0.75; Fig. 2K insert). The overall the concentration of myosin within a cell, unsurprisingly, correlated highly with the concentration in its neighbours. Cells with high concentrations always constricted, and all cells remained inert at low concentrations. But in the range between these values, for any given cell-internal myosin concentration, the cells that expanded were always those for which the neighbours had the highest myosin levels (Fig. 2L, K insert). This shows that forces acting on each cell from its neighbours have an important role in determining the cell’s behaviour.

### Actomyosin model of the mesoderm

We do not know whether the correlation between actomyosin distribution and cell stretching reflects causality, or whether both are effects of external forces, i.e. pulling by neighbours. We used a microscopic, filament-based model^56,57^ to test under what conditions cells containing contractile actomyosin show the behaviours we observe in the embryo. We again used a chain of “membrane”-separated elements with fixed outer boundaries (Fig 2M). Each “cell” contained a constant number of actin filaments and crosslinkers, and membranes had attachment points for filaments. We varied the number of active myosin motors according to the same distributions as in the visco-elastic model.

A set of profiles was able to generate constriction of 6 and stretching of 3 cells, of which a subset reproduced the qualitative behaviour seen *in vivo*, with an inverted pattern of stretching (red region in Fig. 2Q). For such profiles, ‘cell 7’ stretches until its actomyosin network tears apart, disconnecting the more lateral cells from the constricting cells (Fig. 2N-N’). In conclusion, without *a priori* assumptions, this model gives an output consistent with our experimental observations and indicative of non-linear (yielding) behaviour, showing that such behaviour can emerge directly from the properties of the network components and the myosin concentrations. The striking similarity between the parameters of the myosin profiles in the two unrelated models (microscopic and visco-elastic) that yield the same results (though with an offset of 1 cell-width; Fig. 2Q,R) illustrates the generality of the results and suggests that contractile meshworks *in vivo* can, in theory, do the same. Rapid cell expansion due to strain-softening has also been observed in elegant tissue culture experiments, where persistent intermediate filaments allowed re-establishment of connectivity and the cell re-contracting^8^. Our results here are the first demonstration of an equivalent process occurring in a physiological situation *in vivo*.

### Intrinsic versus externally imposed behaviours of mesodermal cells

Our results so far show that cell-intrinsic genetic regulation or myosin levels alone cannot explain the difference between constricting and stretching. We also compared the role of myosin levels among cells at the same position in the gradient. Many central cells expand transiently before constricting and some are internalized without constricting^58^; Suppl. Fig. 6). We tested whether myosin levels correlated with these behaviours by categorising cells from all rows as either ‘transiently expanding’ or ‘contracting’. In rows 3 −5, transiently expanding cells started out with slightly larger surfaces, but the same myosin concentrations as contracting cells. During the transient expansion (150-250 seconds) neither myosin amounts nor concentration are pronouncedly different from the contracting cells (175 sec; Suppl. Fig. 6). Myosin amounts in row 6 (Suppl. Fig. 6F-J) also rose simultaneously in both populations during the expansion period, the slight divergence in concentration therefore coinciding but not preceding expansion. Thus, myosin levels did not predict constriction versus transient expansion. Finally, central cells that remain unconstricted often have highly asymmetric myosin foci (Fig. 2G-H), much like lateral expanding cells, showing that skewed myosin is not determined by the cell’s position in the genetic gradient, nor by its myosin values. Instead, it seems that myosin distribution in stretching cells is a consequence rather than a cause of their apical size. Together these results suggest that whether a cell constricts does not depend primarily on myosin levels, but at least in part on what its neighbours do, and in part by stochastic variation in its actomyosin organization.

We therefore propose a model where all mesodermal cells have the capacity to constrict in principle, but cells that accumulate active actomyosin earlier or at higher levels than neighbouring cells have a greater chance of sustaining their contraction. This hypothesis makes two testable predictions: (a) preventing central cells from constricting early should allow lateral cells to constrict, and (b), making lateral cells constrict early should affect the ability of central cells to constrict.

To test these predictions, we manipulated apical contractility by laser ablation and optogenetic methods. We first inhibited constriction in central cells by laser-mediated severing of the actomyosin meshwork (Fig. 3). This strongly reduced apical constriction in the illuminated area, and some cells in rows 7 and 8 now constricted their apical sides (Fig. 3D-D”, 3G, Suppl. video 4). Optogenetically inactivating the actomyosin meshwork^59^ yielded the same results: constriction in the illuminated cells was inhibited, several cells in rows 7 and 8 constricted (Fig. 3H-K”). Thus, when central cells are prevented from constricting, lateral cells are able to constrict.

**Fig 3.**
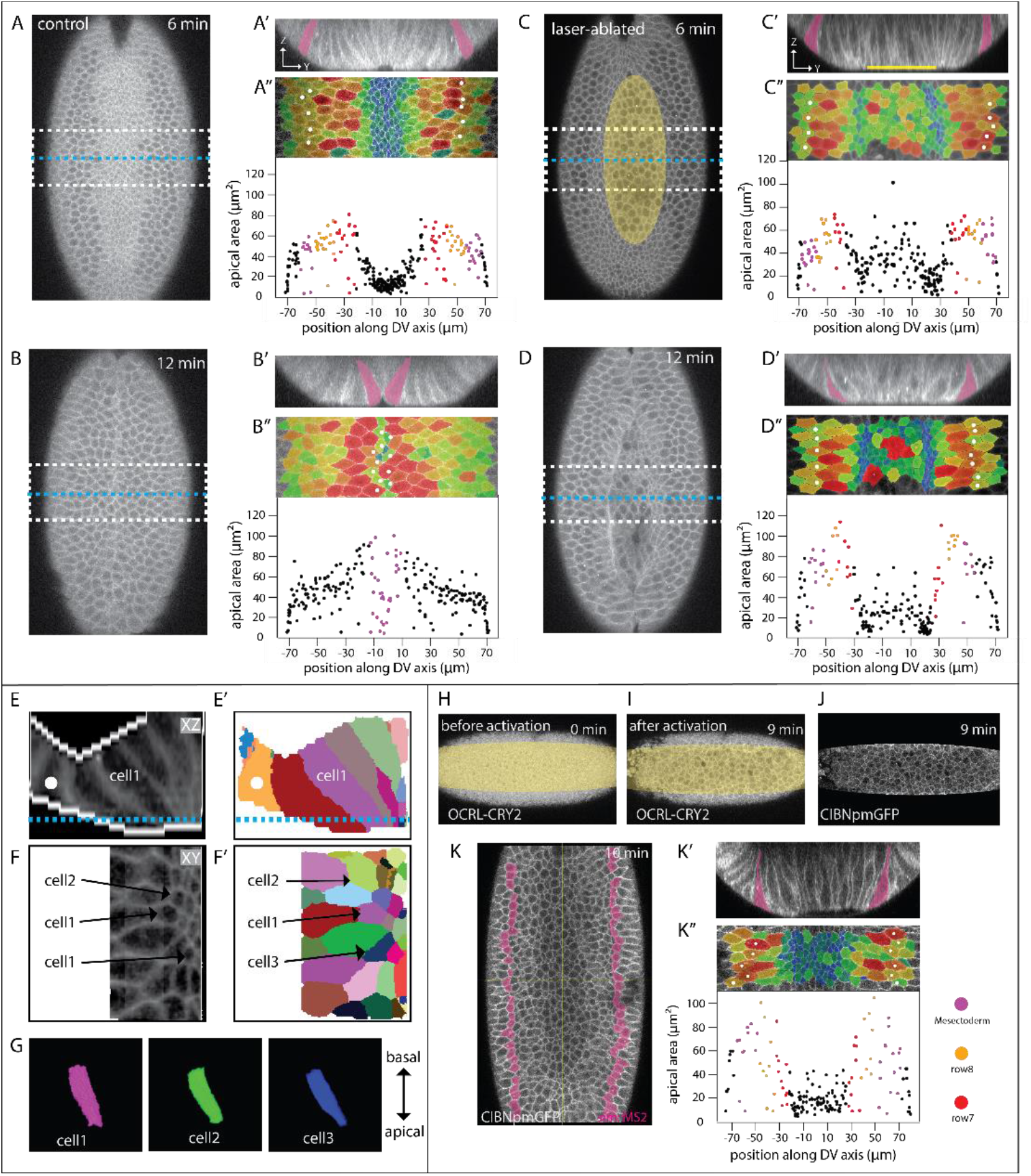
Effects of restricting apical constriction in central cells. (A – D) Two time-points from confocal recordings of control (A, B) and laser-manipulated (C, D) embryos expressing GAP43::mCherry (cell outlines). SnailMS2 and MCP::mCherry (not shown) were used to determine the extent of the mesoderm; mesectoderm is marked by magenta fill in A’ - D” and white spots in A” - D”. 0 sec is the point when the apical-basal length of the central rows is 35μm. The region marked in yellow in (C) was repeatedly illuminated with an infrared laser. See also Suppl. video 3. (A-D) Confocal Z-planes 15μm below the ventral surface. Positions of Z-sections in A’-D’ are marked by yellow lines and the region of the apical surface peels in A”-D” by white boxes. (A’-D’) Z-sections at the positions indicated in A - D. (A” – D”) Apical surface peels of regions marked in A - D. Same markings as in figure 1, with quantification of the apical areas of the cells plotted against their position. Same representation of cell size as in Fig. 1. Note data points at the sides include artefactually small values because cells at the edge are not full size. (E – G) 3D segmentation of l cells from the embryos in (D). (E) Z-section showing cell outlines and the binary mask used for segmentation (white edges). Blue line indicates position of Z-section shown in F. Mesectoderm: white spot. (E’) segmentation result. (F) Z-plane and (F’) segmentation result. The numbers indicate the cells shown in 3D below. (G) 3D renderings of the three cells marked above (F, F’). For 3D viewing see Supplementary movies. See also Suppl. video 4. (H - K) Optogenetic inactivation of cortical actomyosin in a ventrally mounted embryo co-expressing OCRL-CRY2::mCherry, CIBN::pmGFP and simMS2 and MCP::GFP to mark the mesectoderm (H – J) Confocal Z-planes 5 μm below the surface before and after laser-illumination to release actomyosin from the apical cortex. Illumination leads to recruitment of OCRL-CRY2 to the plasma membrane (compare H and I) via membrane-associated CIBN::pmGFP. (K) Z-plane 25 μm below the ventral surface 10 min after laser treatment to show the position of the edge of the mesoderm. Mesectoderm in magenta. This level does not show the apical surface. (K’) Cross-section showing non-stretched cells adjacent to the mesectoderm (magenta). (K”) Apical surface peel and quantification of apical cell areas. Same markings as above.

To test whether the central cells can be stretched, we optogenetically induced premature constriction in lateral cells^10^. We activated regions either side of the central two rows but only in the posterior half of the embryo, retaining the anterior half as control. In the control half, central cells constricted and a gradient of apical areas developed (Fig. 4A’-D’). In the experimental half, ectopic apical constriction occurred in the illuminated cells. At the same time, many of the cells near the ventral midline now expanded their apical surfaces (Fig. 4A”-D’). Thus, central cells failed to undergo their normal morphogenetic programme, even though they themselves had not been manipulated, showing that external forces were able to override their genetic instruction to constrict.

**Fig 4.**
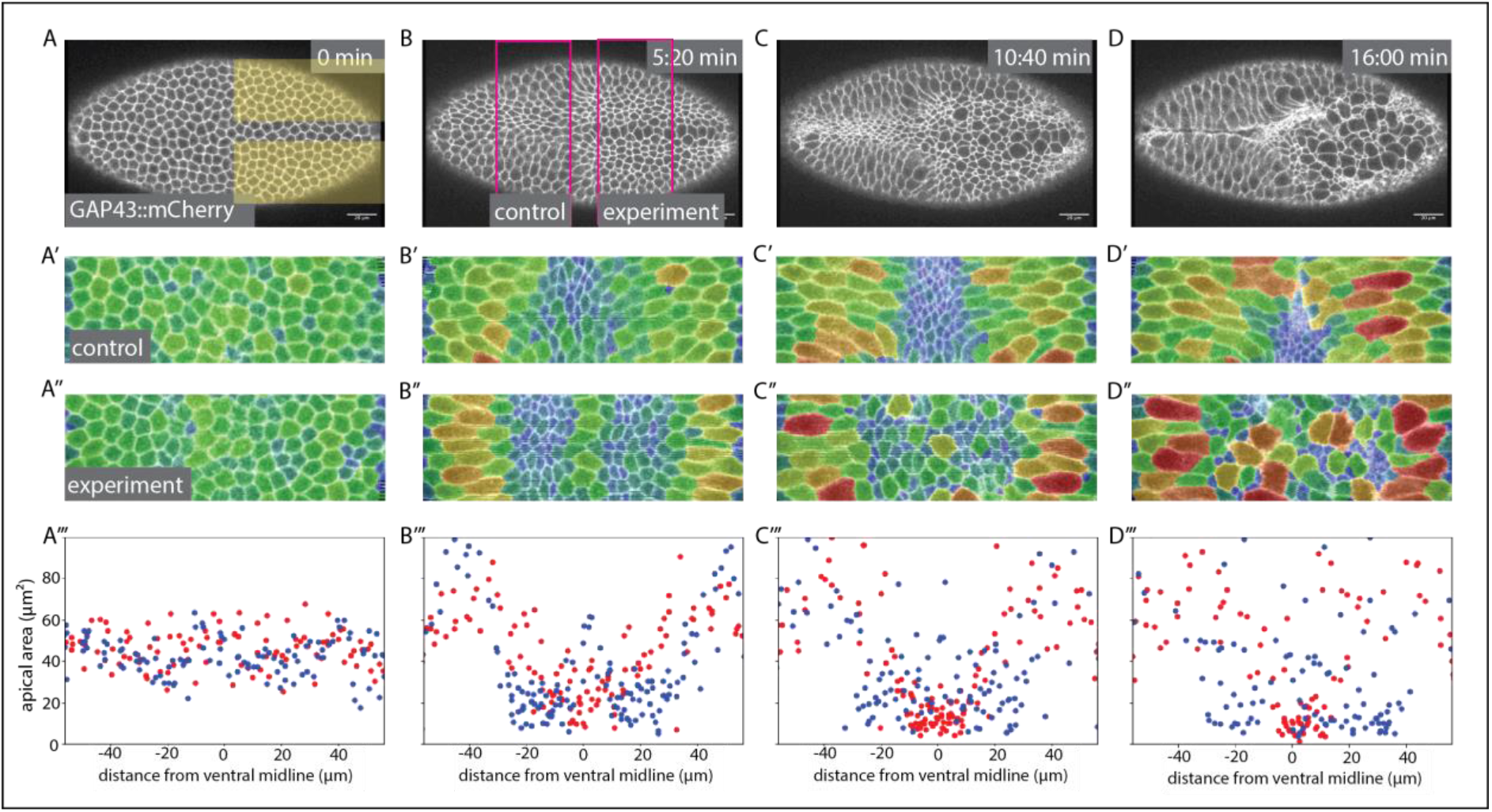
Effect of ectopic myosin recruitment. (A-D) Confocal Z-planes 5μm below the surface of an embryo co-expressing GAP43mCherry, CIBN::pmGFP and RhoGEF2-CRY2. Photoactivation in the yellow areas in (A) induces membranerecruitment of RhoGEF2-CRY2. Magenta lines in (B) show control and experimental areas analysed. (A’-D’, A”-D”) Apical surface peels of the regions marked in (B) overlaid with colour-code representing relative apical areas. (A’”-D’”) Apical areas of the cells in the control (red dots) and experimental (blue dots) parts of the embryo plotted against their positions.

## Discussion

Following from the above, an explanation is needed why lateral cells normally do not constrict, even though they reach sufficient myosin levels. The simplest explanation is that the external forces acting on them are greater than those acting on the early-constricting central cells. While different external forces are likely part of the explanation, in the absence of precise measurements at a subcellular level (an extremely challenging task given the cells’ small size and rapid movement) we must also consider other possibilities.

According to our visco-elastic model, a non-linear stress-strain relation is necessary for the inverted pattern of stretching of lateral cells, which could not be reproduced with previous computational models. The strong stretching, also documented in epithelia *in vitro*^8^, was best recapitulated by a superelastic response. The non-linearity emerging from the microscopic model, however, resembles elastoplasticity (irreversible strain), but the simulations do not include actin turnover which would facilitate recovery from yielding of the cytoskeletal network and thus reverse the stretching, typical of superelastic materials. It is currently not feasible to determine experimentally whether cells in the embryo behave like elastoplastic or superelastic materials (as seen *in vitro*^8^ and simple organisms^53^). Other possible explanations for the same output include dissipation through viscosity^38,63^ or external friction^3,4^, or a non-proportional causal relationship between myosin concentration and constriction forces. The former cannot explain single cell stretching in the central mesoderm, while the latter is unlikely given that myosin levels alone predict a wide range of morphogenetic movements in *Drosophila^51^*

A source of this non-linearity may be the actomyosin not assembling in the proper structure. The pulsatile apico-medial actin meshwork needs to be tightly connected to the junctional complexes to function^13,14,42,60,61^ relying also on an underlying non-pulsatile actin meshwork^62^. Despite the homogeneous actin meshwork in stretching cells, the areas that are free of active myosin occupy a large proportion of the apical surface – similar to ectodermal or amnioserosa cells in which the connection of pulsatile foci to the underlying actin meshwork is lost^62^. The observation that a skewed myosin distribution is not restricted to cells with low myosin but can occur even in central cells at the highest myosin concentrations underscores the conclusion that all aspects of this phenotype are externally imposed rather than intrinsically determined by myosin levels.

Dilution of cortical myosin may compromise the cell’s ability to make sufficient physical connections, in particular along the dorso-ventral axis, so that even if sufficient force is generated, it cannot shorten the cell in the long dimension. In other words, even though the cells have enough myosin to create force, the system is not properly engaged and its force is not transmitted to the cell boundary. In this model, the skewed myosin distribution is both a result of external forces and also part of the cause of a cells’ failure to constrict. By a feed-forward mechanism, an initial expansion induced by constricting neighbours dilutes or distorts the apical actomyosin, giving these cells a lower chance of generating or sustaining a contraction. This mechanism, which we propose corresponds to the non-linear behaviour predicted by the models, would apply both to central and to lateral cells, with a catastrophic ‘flip’ being stochastic and rare in central cells, but reproducible in lateral cells because of the temporal and spatial gradient in which contractions occur.

## Acknowledgments

We thank Mayank Kumar and Catarina Carmo for help with generating fly lines; Dimitri Kromm and Lars Hufnagel for expert help for MuVi SPIM imaging; Marvin Albert for support on image registration; Stefano De Renzis, Hernan Garcia, Thomas Lecuit for stocks and reagents; the EMBL Advanced Light Microscopy Facility (ALMF) for continuous support; Alexandre Cunha and Thiago Vallin-Spina for providing access to SEGMENT3D; Steffen Lemke, Karen Daniels, Justin Crocker, Stefano de Renzis, Aissam Ikmi, Xavier Trepat, Pavel Tomancak and the Leptin lab for critical comments and discussions. This work was supported by funding from EMBO and DFG grant FOR1756.

## Author Contributions

Conceptualization, S.B., M.L.; Methodology and investigation, S.B., J.S., D.G., G.M. V.Z.; Formal Analysis, S.B., D.G., G.M., V.Z.; Writing – Original Draft, S.B., J.M.B, M.L.; Writing – Review & Editing, S.B., M.L.; Visualization, S.B., J.S., D.G., V.Z.; Simulations, J.M.B, G.M.; Supervision, A.K, J. M. B.; M.L.; Funding Acquisition, M.L.

## Supplementary information

### Plasmid for membrane-associated mCardinal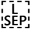

To generate the plasmid attb-tubulin_promoter-GAP43::mCardinal-K10 plasmid, the attb-UASp-K10 plasmid (provided by Anne Ephrussi, EMBL Heidelberg) was modified by replacing the UAS promoter by a tubulin promoter sequence that was amplified from the plasmid pCasper4-tubulin (provided by Stefano De Renzis, EMBL Heidelberg). The mCardinal coding sequence was amplified from mCardinal-H2B-C-10 (Addgene plasmid #56162) using a forward primer with the sequence encoding the first 20 amino acids of the GAP43 protein from Bos taurus (Table S6). The GAP43::mCardinal fragment was inserted into the attb-tubulin-promoter-K10 plasmid using NotI and BamH enzymes.

### Generation of fly stocks

To generate the fly transgenic lines p[mat tub>GAP43::mCardinal]/Cyo and p[mat tub> GAP43::mCardinal]/TM6 Tb, the attb-tubulin_promoter-GAP43::mCardinal-K10 plasmid was inserted into landing sites on the second and third chromosomes (landing sites VK18 (#BDSC-9736) and VK33(#BDSC-9750)) by BestGene Inc. (California, USA). Only the insertion on the second chromosome was used in this study because it was brighter than the insertion in VK33.

### Sample preparation

Embryos were collected according to standard procedures on apple juice agar plates. Plates were changed after a one-hour embryo collection and kept at 25°C for 2.5 hours. Individual mid-to-late cellularization embryos were hand-selected under halocarbon 27 oil. The stage-selected embryos were devitellinised with 50% bleach and washed thoroughly with distilled water. For confocal microscopy, the embryos were then mounted on a glass-bottom microwell dish with the ventral or ventral-lateral side facing the glass and covered with PBS. For MuVi-SPIM the embryos were mounted in 1% Gelrite inside a glass capillary and multiple views registered and fused^16^.

### Confocal microscopy

For visualising 3D cell shapes using 2-photon illumination, a femtosecond-pulsed infrared laser (Chameleon Compact OPO Family, Coherent) tuned at 950 nm emission wavelength and coupled with Zeiss LSM 780 confocal microscope was used. The region of interest was defined with the Zen ‘Regions’ interface and the embryos were illuminated with 20-25% laser power. A volume of 200 x 500 x 60 μm^3^ was imaged, where the dimension of 200 μm is along the anterior-posterior axis of the embryo, centred around the central region, 500 μm is along the left-right axis, and 60 μm is depth in the z axis.

Two-colour imaging was performed at room temperature with a Zeiss 880 Airyscan microscope, a 40X/1.4 numerical aperture oil-immersion objective, an argon ion laser and a 561-nm diode laser. Image stacks were acquired every 25 sec.

### Selective plane illumination microscopy

Imaging was performed on a custom-built Multi-View SPIM set-up^68^ with Nikon 10/0.3W objective lenses for illumination and Nikon 20/1.0W objective lenses for detection. An additional 1.5X magnification tube lens produced an effective image pixel size of 0.19 μm X 0.19 μm. Optical sections were recorded with a typical spacing of 0.75-1 μm. For observing cell shape changes, *GAP43::mCardinal* embryos were imaged from two opposing directions simultaneously and successively from two directions with 90 degree apart. Registration of the four views was performed as previously described^16^.

### Identification of mesodermal cells

To identify unambiguously the lateral borders of the mesoderm we used two methods (Suppl Fig. 1D-J’): i) back-tracing mesectodermal cells from the point when the meet at the ventral midline after the mesoderm is fully internalised; or ii) using the MS2 stem loop/MCP-GFP system to visualize the expression of the genes *singleminded*^69^ in mesectodermal cells or *snail* in the mesoderm^70^.

### Laser ablation and illumination

Laser-based actomyosin meshwork ablation was performed as previously described^16^ using a femtosecond-pulsed infrared laser (Chameleon Compact OPO Family, Coherent) tuned at 950 nm emission wavelength and coupled to an LSM Zeiss 780 confocal microscope. The Zen ‘Bleaching’ interface was used to create the region of interest and was illuminated at 65-70% laser power. For this experiment, C-Apochromat 63X magnification water immersion Zeiss Objective with 1.1 NA was used (infrared corrected).

### Optogenetic manipulations

Embryos were prepared in a room where the blue spectrum of visible light was filtered out^10^. The Zen ‘Regions’ interface was used to create the region of interest and the embryos were illuminated with 15-20% laser power with pixel dwell time between 0.8 and 1.27 ms. For this experiment, a C-Apochromat 40X magnification water immersion Zeiss Objective with 1.2 NA was used (infrared corrected) and an infrared laser (Chameleon Compact OPO Family, Coherent) tuned to 950 nm emission were used.

### Image processing

#### Apical surface extraction from SPIM images

The 30% of central part the embryos along the anterior-posterior axis was cropped in Fiji^71^. A custom MATLAB software was then used to extract the apical surfaces^36^. A binary mask around the embryo was generated semi-automatically by defining the apical and basal surfaces. Using these masks, distance transformation was used to define a 1 to 2-pixel ‘peel’ typically 2-3 pixels below the binary mask. Along the anterior-posterior axis of the embryo, pixels along the surface were traced and mapped onto a line. This process was performed on every stack to map the apical surface of the embryo onto a 2D plane.

#### Myosin measurements

Images were deconvolved in the ZEN software using AiryProcessing. The Spider:GFP images represent confocal slices 3 μm below the apical cortex. Sqh:Cherry images represent sum *Z*-projections of an apical section of the same depth upon background myosin subtraction. Background myosin intensity was measured in single subapical confocal slices, mean + 2 standard deviations were subtracted from each slice before Z-projecting to obtain apical myosin intensity. The cells were segmented and tracked using TissueAnalyzer^72^. The segmentation output was used to extract cell areas and pixel intensities. Myosin intensity within a cell was measured as a sum intensity of all pixels in a cell. Myosin concentration was calculated as myosin intensity/cell area.

For Figures 2K and 2L, myosin concentration and cell size values for every cell were smoothed along the time axis using a 1D gaussian filter with sigma=3 (reference^73^). For every cell at each of 25 time points three values were taken: its myosin concentration, the myosin concentration in the area of 70 pixels around the cell boundary and the relative size change, calculated as the cell size in the next time frame divided by the cell size in the current one.

#### A visco-elastic model for the mesoderm

We modelled the mesoderm as one-dimensional series of points (cell boundaries) connected by visco-elastic units (cells). Each cell behaves as a Kelvin-Voigt material made of a spring and dashpot in parallel connecting two adjacent cell boundaries (at positions *x_i_* and *x*_*i*+1_). All cells have the same viscosity (*η*) and stress-strain response (*S*(Δ*x*)). We added 3 cells with a higher stiffness at each side of the 19 mesodermal cells to simulate the rigid ectodermal cells. Each cell contains a defined amount of “myosin” (*M*), which exerts a force at each cell membrane position *x_i_* that is directly proportional to the local gradient of “myosin” around that point (*∇M*(*x_i_*)). The system evolves over time according to the following deterministic equation:

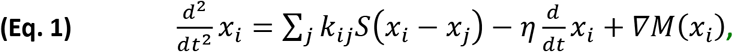

where the sum is over the two adjacent point coordinates, the function *S* is the stress-strain response (defined below), and the myosin profile (*M*) is modelled as a symmetric sigmoidal function around the midline, described by the equation:

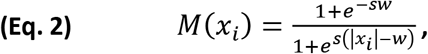

where *w* and *s* are parameters describing the width and steepness of the function, respectively, and *x_i_* = 0 corresponds to the midline (central) position of the mesoderm.

We considered 5 types of stress-strain responses models. The first corresponds to a simple linear elastic-like model, where stress increases proportionally with the strain. The other four models are non-linear, with the same stiffening response to compressive strains (Δ*x* < 0), and 4 different types of responses to extensive strains: i) an elastomer-like model, corresponding to an elastic response but with a decreased stiffness after the proportionality limit; ii) a stiffening model, with an increased stiffness after the proportionally limit; iii) a superelastic model, corresponding to a material that undergoes strain-softening after the proportionality limit, followed by strain-hardening; and iv) an elastoplastic model, with a similar curve as before, but undergoing plastic (permanent) deformation after a certain yielding stress. For simplicity, all stress-strain curves are continuous functions, with a repulsive response for compressive strains (Δ*x* < 0) to prevent cells from having zero areas and made of connected linear segments with varying slopes (stiffness) for different ranges of extensive strains (Δ*x* > 0). Table S1 contains a mathematical description of each curve, and Table S2 lists the parameters values used in our simulations.

We systematically explored the outcomes of the 5 models by varying the parameters controlling the myosin profile (Eq. 2).

#### Microscopic model for the mesoderm

We modelled a line of cells from the ventral midline to the mesectodermal cell as a series of sequentially connected actomyosin networks with varying amounts of myosin motors. Each network is a 2D mesh of 800 actin filaments of 1.5 μm long, randomly distributed within a rectangular region of 7 x 8 μm. The cells have periodic boundary conditions along the “antero-posterior” direction (top to bottom in the graphic representation) and are separate by rigid but movable ‘membranes’. The row is bounded by unmovable walls on each end to simulate the ectoderm and the ventral midline. Each membrane has 800 connecting points for the filaments on each side. Actin filaments of adjacent cells do not interact except through the membrane connectors. Each cell has 1,600 crosslinkers and between 1,600 to 16,000 myosin motors (with a minimum level that was sufficient in principle to contract the network). Both connectors are modelled as point like objects with two independent hands that can bind and bridge two nearby filaments pertaining to the same cell. Once bound, motor hands move towards the plus-end of the filaments until they unbind or reach and detach from their ends. We used as input the parameters for the crosslinkers, connectors, motors and filaments the on-off rates, movement kinetics and stiffness/persistence lengths that have been biochemically determined for alpha-actinin, myosin and F-actin (see Table S3).

All filament-based simulations were done with CytoSim^57^, a cross-platform simulation engine designed to handle large systems of flexible filaments and associated proteins. CytoSim uses a Brownian dynamics approach to simulate the cytoskeleton, where each element is individually represented in either 2D or 3D space. The number, spatial location and physical properties of each element is determined at the start of the simulation and the system evolves according to the laws of mechanics and stochastic reaction-kinetics.

### Data Analysis and plotting

All graphs were plotted using either MATLAB (MATLAB_R2015a) or Python (version 3.6). Matplotlib^78^, Pandas^79^, Scikit-image^80^, NumPy^81^ packages were used.The figures were compiled using Adobe Illustrator CS6 (Version 16.0.0).

### Data availability

Apart from the third party software tool SEGMENT3D^74^, all described algorithms were implemented in MATLAB and are available from https://github.com/stegmaierj/CellShapeAnalysis/ (Apache License 2.0) and the code for myosin analysis from https://github.com/sourabh-bhide/tissue2cells

**Table S1:** Description of equations used for the visco-elastic stress-strain responses.

| Stress-strain type | Strain range |  |  |  |
| --- | --- | --- | --- | --- |
| | $\Delta x < 0$ | $\Delta x < x_{pl}$ | $\Delta x < x_{sh}$ | $x_{sh} < \Delta x$ |
|  | function | slope | slope | slope |
| Linear | $\Delta x s_1$ | $s_1$ | $s_1$ | $s_1$ |
| Elastomeric | $k_c [x_c^{-n} - (\Delta x + x_c)^{-n}]$ | $s_1$ | $s_{2em}$ | $s_{2em}$ |
| Stiffening | $k_c [x_c^{-n} - (\Delta x + x_c)^{-n}]$ | $s_1$ | $s_{2s}$ | $s_{2s}$ |
| Superelastic | $k_c [x_c^{-n} - (\Delta x + x_c)^{-n}]$ | $s_1$ | $s_{2se}$ | $s_{3se}$ |
| Elastoplastic | $k_c [x_c^{-n} - (\Delta x + x_c)^{-n}]$ | $s_1$ | $s_{2ep}$ | $s_{3ep}$ |

**Table S2:** Parameters used in the visco-elastic models.

| Parameter | Value |
| --- | --- |
| $k_c$ | 131.247 |
| $x_c$ | 6 |
| $n$ | 0.05 |
| $s_1$ | 1.0 |
| $s_{2em}$ | 0.2 |
| $s_{2s}$ | 2.0 |
| $s_{2se}$ | -0.75 |
| $s_{3se}$ | 0.2 |
| $s_{2ep}$ | -1.25 |
| $s_{3ep}$ | 0.2 |

**Table S3:** List of parameters used in the microscopic simulations.

| Parameter | Symbol | Value (or range) | Units |
| --- | --- | --- | --- |
| Simulation time step | $\Delta t$ | 0.01 | s |
| total time | T | 20 | s |
| viscosity | $\eta$ | 0.1 | pN s/$\mu\text{m}^2$ |
| Geometry |  |  |  |
| cell height | H | 8 | $\mu\text{m}$ |
| cell width | W | 7 | $\mu\text{m}$ |
| number of cells | $N_c$ | 9 | - |
| Actin filaments |  |  |  |
| length | L | 1.5 | $\mu\text{m}$ |
| rigidity | k | 0.0075 | pN $\mu\text{m}^2$ |
| segmentation | $\Delta L$ | 0.1 | $\mu\text{m}$ |
| filaments per cell | $N_F$ | 800 | - |
| Myosin motor |  |  |  |
| binding rate | $k_{mb}$ | 10 | $\text{s}^{-1}$ |
| binding range | $r_{mb}$ | 0.1 | $\mu\text{m}$ |
| unbinding rate | $k_{mu}$ | 0.3 | $\text{s}^{-1}$ |
| unloaded speed | $v_0$ | 0.1 | $\mu\text{m/s}$ |
| stall force | $f_0$ | 6 | pN |
| stiffness | $s_m$ | 500 | pN/$\mu\text{m}$ |
| motors per cell | $N_m$ | 1600-16000 | - |
| Crosslinker |  |  |  |
| binding rate | $k_{xb}$ | 10 | $s^{-1}$ |
| binding range | $r_{xb}$ | 0.1 | $\mu m$ |
| unbinding rate | $k_{xu}$ | 0.3 | $s^{-1}$ |
| stiffness | $S_x$ | 500 | $pN/\mu m$ |
| crosslinkers per cell | $N_x$ | 1600 | - |
| Membrane connectors |  |  |  |
| binding rate | $k_{cb}$ | 2000 | $s^{-1}$ |
| binding range | $r_{cb}$ | 0.1 | $\mu m$ |
| unbinding rate | $k_{cu}$ | 0 | $s^{-1}$ |
| stiffness | $S_c$ | 500 | $pN$ |
| connectors per membrane | $N_{cm}$ | 400 | - |
| connectors per wall | $N_{cw}$ | 800 | - |

**Table S4:** List of Fly Stocks.

| Stock | source | Reference |
| --- | --- | --- |
| w[*]; p[UAS-sqh-Gap43::mCherry]/CyO; + | Thomas Lecuit | 13 |
| w[*]; p[sqh-MRLC::eGFP]/CyO; p[UASp-Gap43::mCherry]/MKRS | Thomas Lecuit (sqh-GFP) | 75 |
| sqh<sup>AX3</sup>; p[sqh-UtrophinABD::GFP], p[sqh-MRLC::mCherry] | Thomas Lecuit (sqh-mCherry) | 12 |
| sqh<sup>AX3</sup>; p[sqh-MRLC::mCherry]; p[Spider::GFP] | Stefano DeRenzis | 13 |
| w[*]; p[snail::MS2]; + | Jacques Bothma | 70 |
| w[*]; p[MCP::mCherry]/CyO ;<br>p[MCP::mCherry]/TM3,Ser | Jacques Bothma | 76 |
| w[*]; + ; P[w+,UASp-mCherry::CRY2- OCRL]/Sb | Stefano DeRenzis | 59 |
| w[*]; P[w+,UASp-CIBN::pmGFP]//CyO ;<br>sb/TM3,Sb | Stefano DeRenzis | 59 |
| w[*]; p[MCP::GFP] | Stefano DeRenzis | 77 |
| w[*]; p[Sim::MS2] | Stefano DeRenzis | 69 |
| w[*]; p[UASp-RhoGEF2-CRY2]/TM3, Ser | Stefano DeRenzis | 10 |
| p[sqh::GFP];p[w+,mataαTub-Gal4::VP16],p[UASp-Gap43::mCherry::mCherry]/TM3 | Adam Martin | 49 |
| w[*]; If/CyO; p[Oskp-Gal4::VP16]/TM3, Ser | Bloomington stock 23651 |  |
| p[sqh>Gap43::mCherry]; +; + | Stefano DeRenzis | 10 |

**Table S5:** Materials.

| Product name | Product information |
| --- | --- |
| Glass bottom plates | Matek corporation (Part no.: P35G-1.5-10.C) |
| Microspheres | TetraSpeck <sup>TM</sup> Fluorescent Microspheres, ThermoFisher (Catalogue no.:T7284) |
| Gelrite | Merck (Catalogue no.:G1910) |
| Halocarbon Oil 27 | Merck (Catalogue no.:H8773) |

**Table S6:**
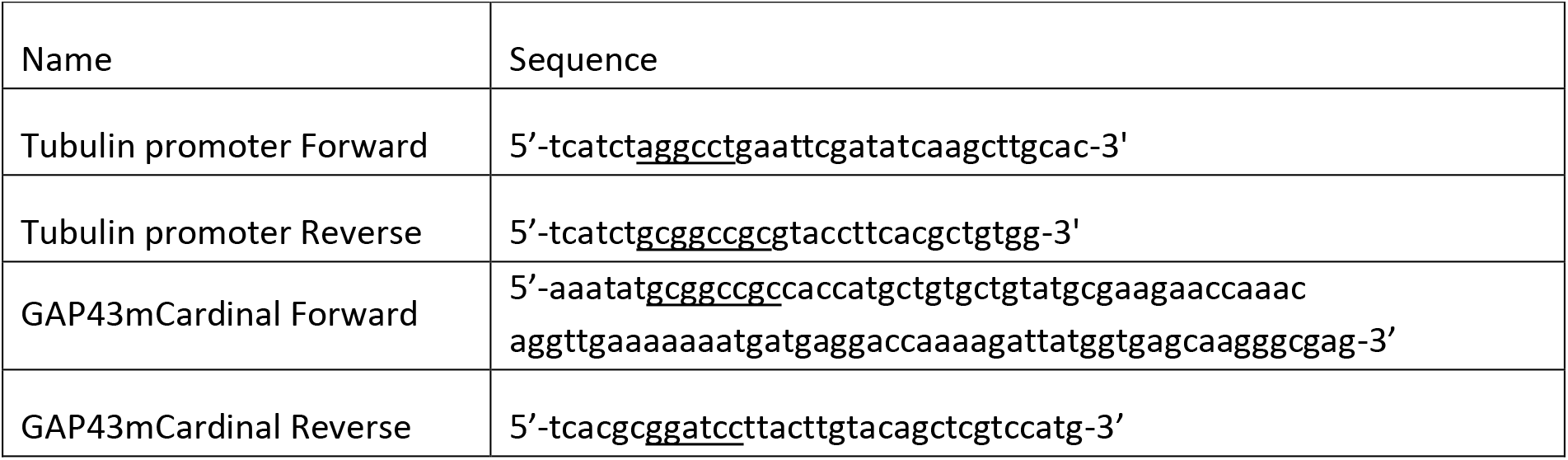
Primers.

**Table S7:** List of genotypes used in experiments.

| Figure no. | Fly stock/ Cross | Microscopy |
| --- | --- | --- |
| 1 C-C'', D-D''; S1 A-C | p[mat tub >GAP43::mCardinal]/CyO | MuVi SPIM |
| 1 E,E'; 2 E-G | sqh<sup>AX3</sup>; p[sqh-MRLC::mCherry]; p[Spider::GFP] | LSM 880 |
| 1 I,I'; S2; S3 | sqh<sup>AX3</sup>; p[sqh-UtrophinABD::GFP], p[sqh-MRLC::mCherry] | LSM 880 NLO |
| 3 A-B , C-D, S1 D-I | sqh-MRLC::eGFP/MCP::mCherry; UASp-Gap43::mCherry/MCP::mCherry X Snail MS2/SnailMS2 | LSM 780 NLO |
| 3 H-K | CIBNpm::GFP/MCP::GFP; OCRLCRY2::mCherry/osk Gal4 X Sim-MS2/Sim-MS2 | LSM 780 NLO |
| 4 | sqhp-Gap43::mCherry/+; UASp>CIBN::pmGFP ; UASp>RhoGEF2-CRY2 / Osk>Gal4::VP16 | LSM 780 NLO |
| S2 A-D | sqh<sup>AX3</sup>; p[sqh-UtrophinABD::GFP], p[sqh-MRLC::mCherry] | MuVi SPIM |
| S6 | sqh<sup>AX3</sup>; p[sqh-MRLC::mCherry]; p[Spider::GFP] | LSM 880 |

## Supplementary figures

**Suppl. Fig. 1.**
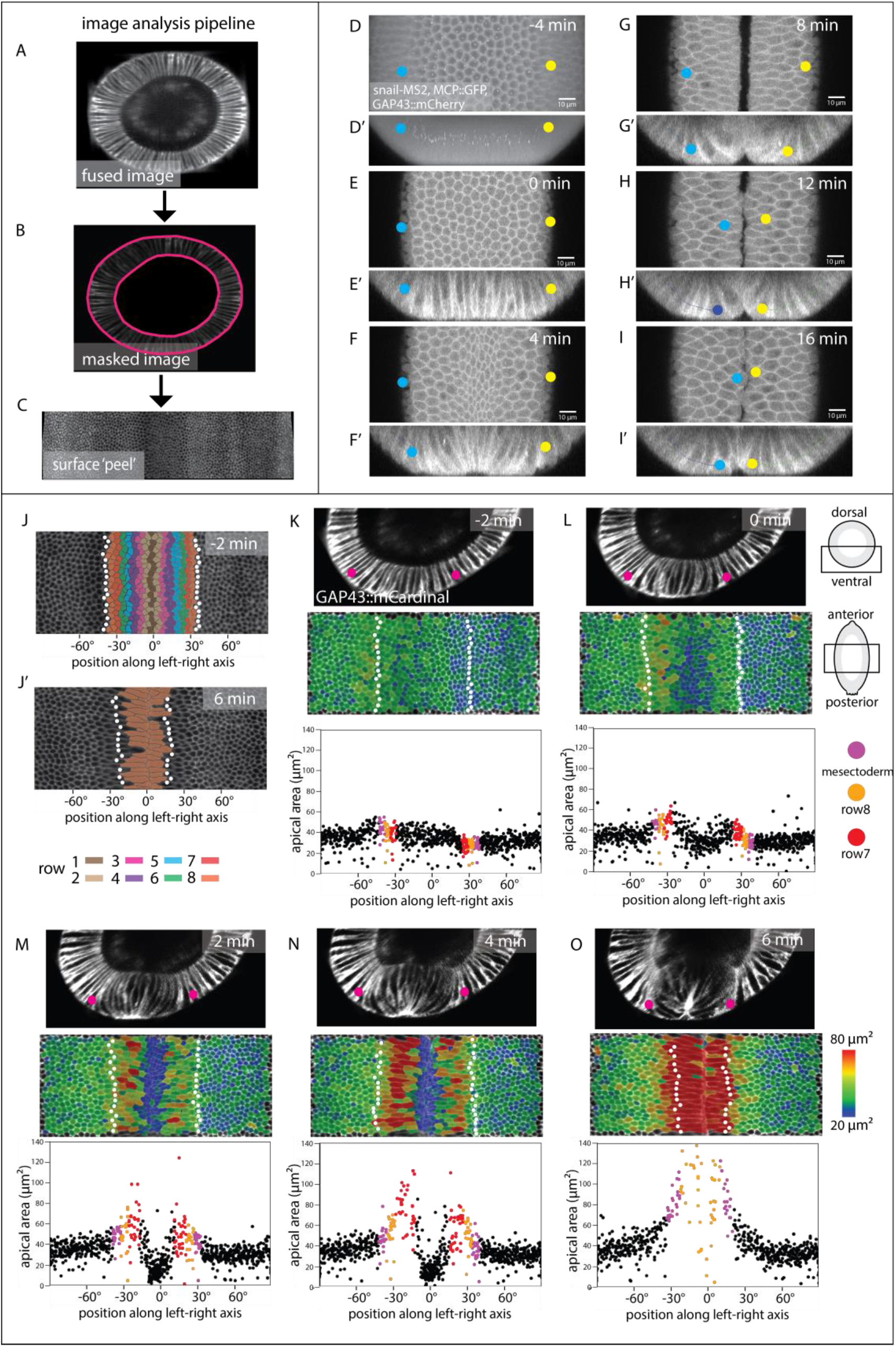
Image analysis and identification of the edge of the mesoderm. (A) Embryos were imaged using Multi-View SPIM. The resulting data sets were fused into a single 3D stack for each timepoint. Ventral side top left. (B, C) Apical (outside) and basal surface masks were defined semi-automatically. These masks were then used to extract the apical surface of the embryo^36^. (D, D’) Maximum intensity projections along the apical-basal (D) and anterior-posterior (D’) direction of an embryo co-expressing GAP43::mCherry, Snail::MS2 and MCP::mCherry 4 min before the initiation of ventral furrow formation. The white spots represent sites of *snail::MS2* RNA in the nuclei of mesodermal cells. Yellow and blue dots mark the positions of the adjacent mesectodermal cell rows. (E-I) Confocal Z-plane 2μm from the surface and Z-sections (E’-I’) over the course of furrow invagination. The mesectodermal cell rows meet at the midline. Back-tracing from this time point can be used to determine the edge of the mesoderm in unmarked embryos. (J, J’) Surface peels extracted from MuVi SPIM images at −2 and 6 min from initiation of ventral furrow formation. White dots indicate the mesectodermal cells as determined by backtracing. Cell rows are colour-coded with numbering coordinated operationally around row 6, which is the last non-stretching row and easily identifiable in all movies, regardless of imaging angle. The width of the mesoderm varies along the anterior-posterior axis, with a width of less than 18 cells in some areas. (K-O) Each panel is from one time-point from a MuVi SPIM recording of an embryo expressing GAP43::mCardinal, with three of the three images showing first, cross-sectional views; secondly, apical surface ‘peels’ extracted from the ventral half of the central one third of the embryo, And finally and third, the apical areas plotted against cell position along the left-right axis (the centre, 0°, is the ventral midline of the embryo). Each dot represents one cell. Apical cell areas measured from segmented images were colour-coded and overlayed on the original image. Mesectodermal cells are marked as white dots in the surface peels and as magenta dots in the plots. For the description in this figure, we define t= 0 min as the time when cells in the central four rows have constricted on average by at least 20%.

**Suppl. Fig. 2.**
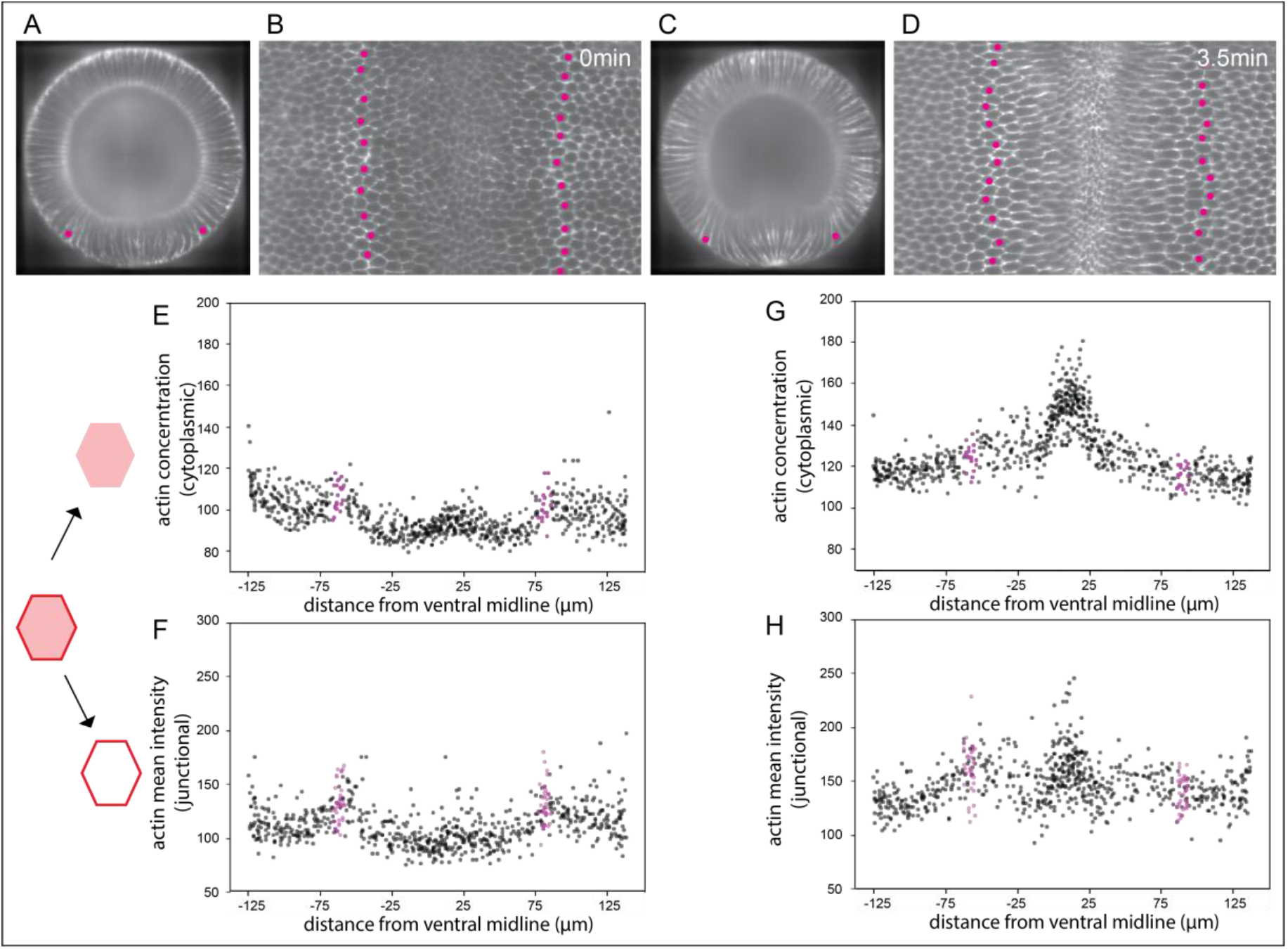
Distribution of apical F-actin in the mesoderm. (A-H) Data from MuVi SPIM recordings of an embryo expressing UtrABD::GFP to visualize F-actin. The rows of mesectodermal cells (identified by back-tracing) are marked in magenta (A,C) Cross sectional view at ~50% egg length and (B,D) sub-apical peel extracted 2μm below the apical surface. (E-H) From the segmented subapical peel, the F-actin mean cytoplasmic concentration (sum of all pixel intensities divided by area) and mean junctional intensity (sum of all pixel intensities divided by length of the junction) are plotted for every cell in the area shown. The mean cytoplasmic intensity and the mean junctional intensity are lower in the mesodermal cells than the ectodermal cells at the onset of furrow formation and increase slightly as furrow formation proceeds.

**Suppl. Fig. 3.**
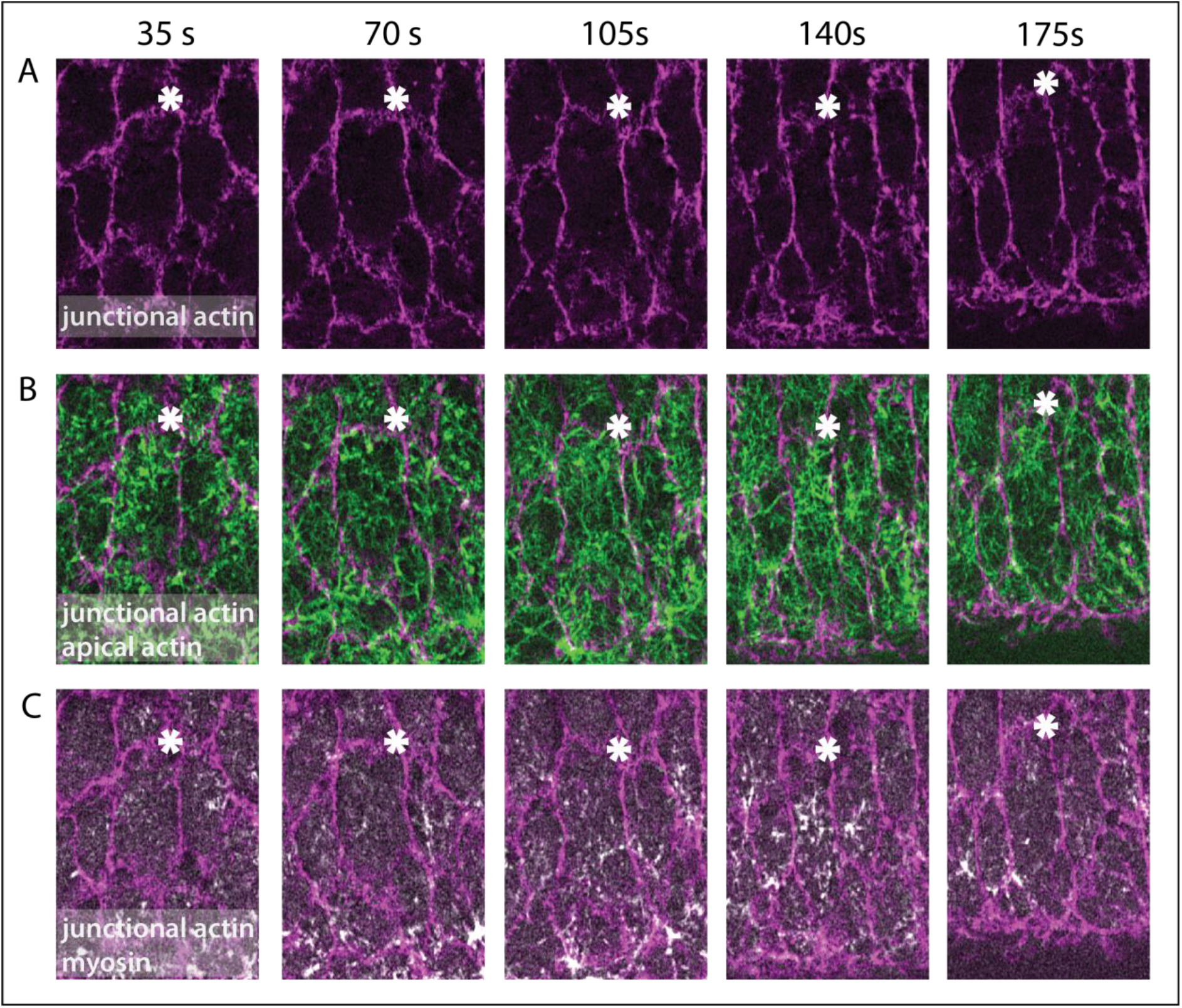
Apical actomyosin meshwork in an expanding lateral cell. Ventro-laterally mounted embryo (same as shown in Fig. 1I-I’) expressing utrABD::GFP and sqh::mCherry to visualise F-actin and myosin. (A) junctional actin in a confocal section 3 μm from the surface. (B) apical cortical actin meshwork (green; sum intensity Z-projection of confocal sections within 1 μm from surface) and subapical junctional actin (magenta) to visualize cell boundaries. (C) sum intensity Z-projections of confocal sections within 1 μm from surface for apical myosin (red) junctional subapical junctional actin (magenta) to visualize cell boundaries. The white asterisk is a reference point.

**Suppl. Fig. 4.**
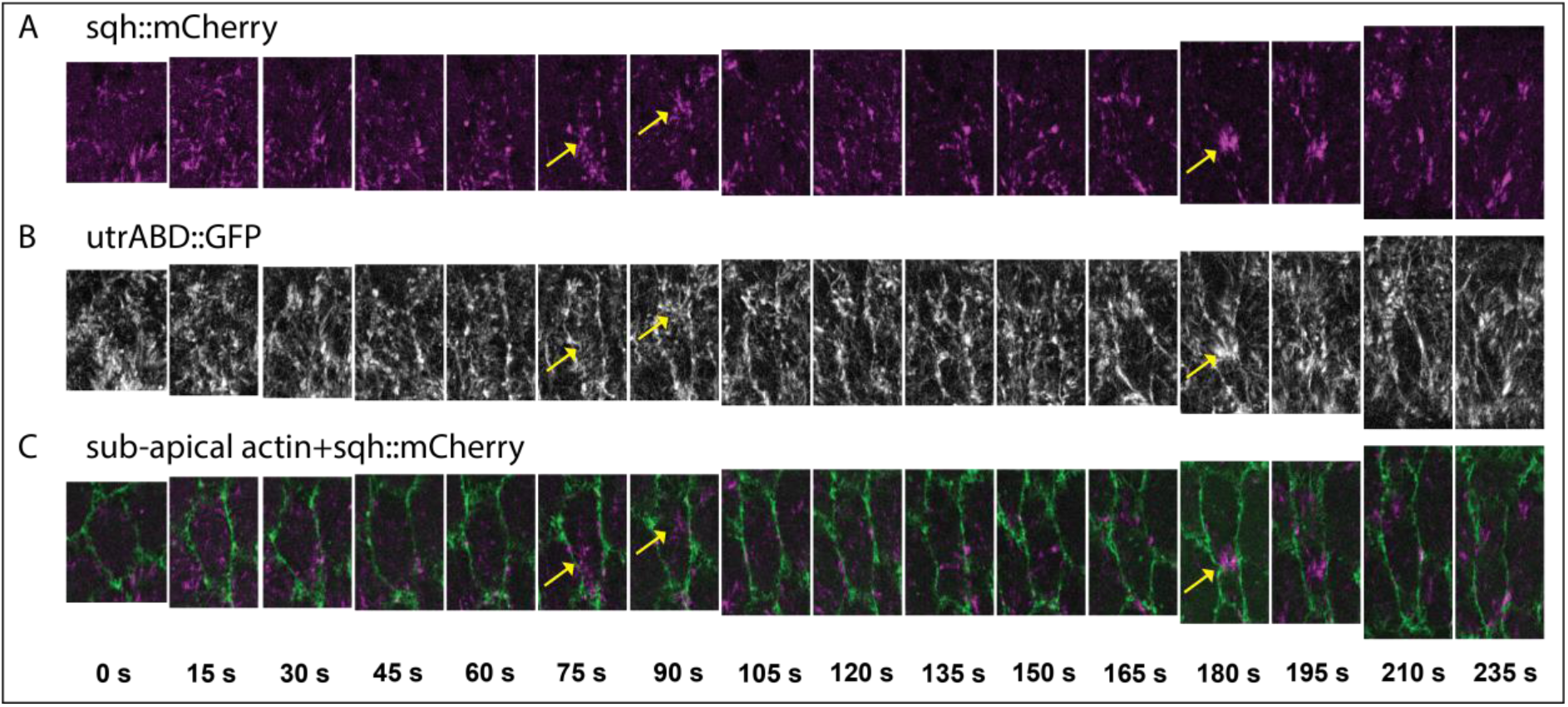
Pulsatile actomyosin meshwork in lateral mesodermal cell. Example of a stretching lateral mesodermal cell in a ventro-laterally mounted embryo (same as shown in Fig. 1I-I’) expressing UtrABD::GFP (white) and sqh::mCherry (magenta). (A, B) Sum intensity projections of myosin (A) and apical actin (B) in the first 2 μm below the surface. (C) Subapical cortical F-actin (2.5 μm below the surface; green) marks the cell boundaries. Actomyosin foci form twice (yellow arrows, 75sec and 180 sec) and create a constriction in the anterior-posterior direction.

**Suppl. Fig. 5.**
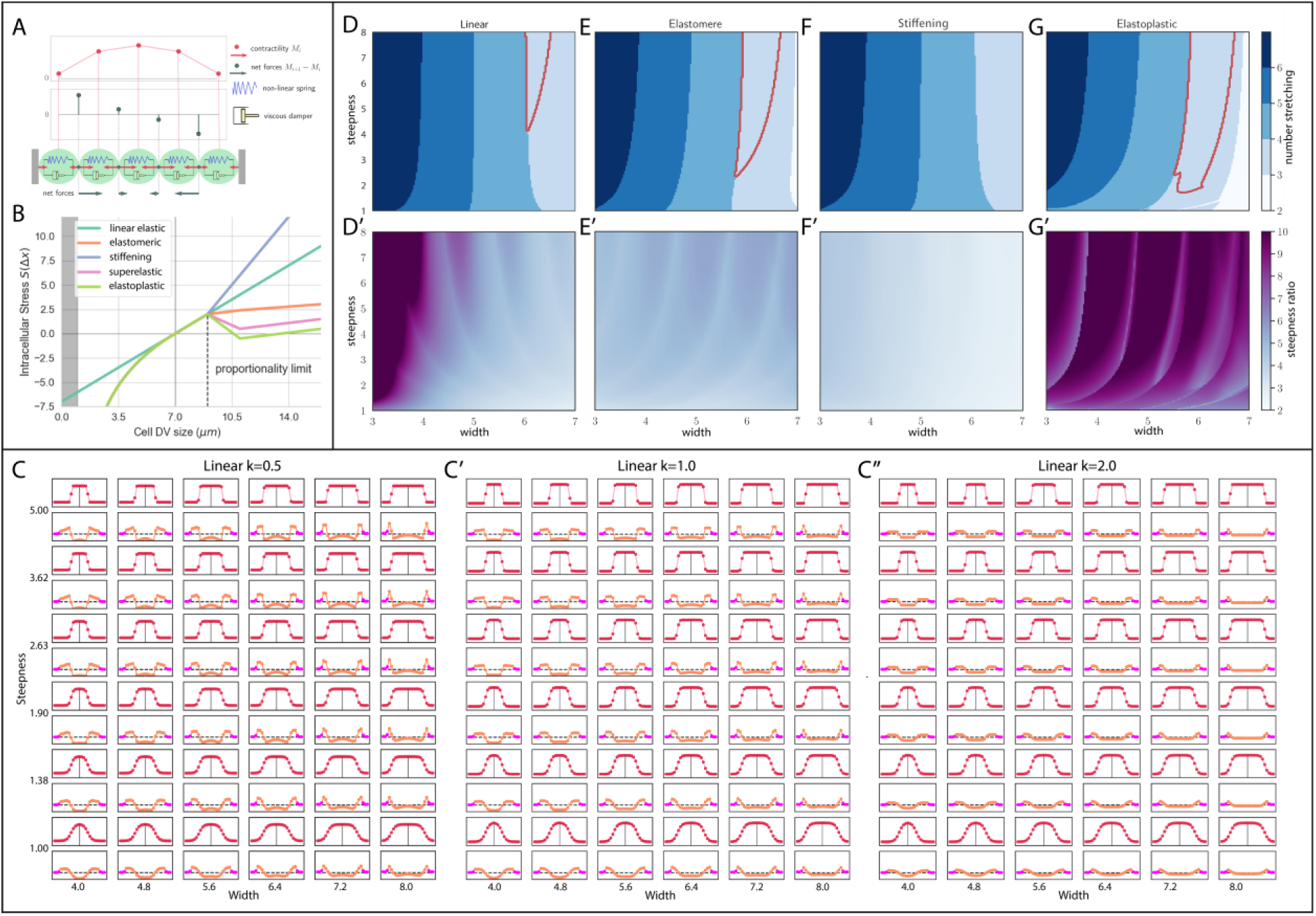
Model of a line of visco-elastic elements representing on line of cells across the mesoderm. (A) Cells are modeled as a series of Kelvin-Voigt viscoelastic elements with viscosity (*η*) and spring constant (*k*). Cell size changes depend on the contractile forces within cells (red arrows ‘pulling’ on connection points) and movement of the connecting points which is determined by the differential forces (grey arrows) acting on them. (B) Graphs for four stress-strain relationships (linear elastic, elastomeric, superelastic and elastoplastic) that are imposed on the spring constants in the model. The resting length of the cell is set to L=7μm. Deviation from the resting length causes either expansion (positive stress) or constriction (negative stress). (C-C”) Parameter scan of the myosin profile with varying steepnesses and peak widths. The myosin profiles (M(x)) are shown in red, and the resulting final cell sizes below in orange. The stiffer ‘ectodermal’ cells are marked in pink. The visco-elastic elements have linear elasticity with constant (k). (D – F, D’-F’) Same marking as Fig. 2O. Parameter map for myosin concentration curves with varying peak widths and steepnesses for linear elastic, elastomeric and elastoplastic materials. Blue shades: number of expanding cells. Red outline: conditions where the three right cells expand with an inverted pattern of stretching that qualitatively matches experiments. Shading in D’ - F’: ratio of the most constricted to the most expanded cell; magenta: largest size differences, light blue: minimal size differences.

**Suppl. Fig. 6.**
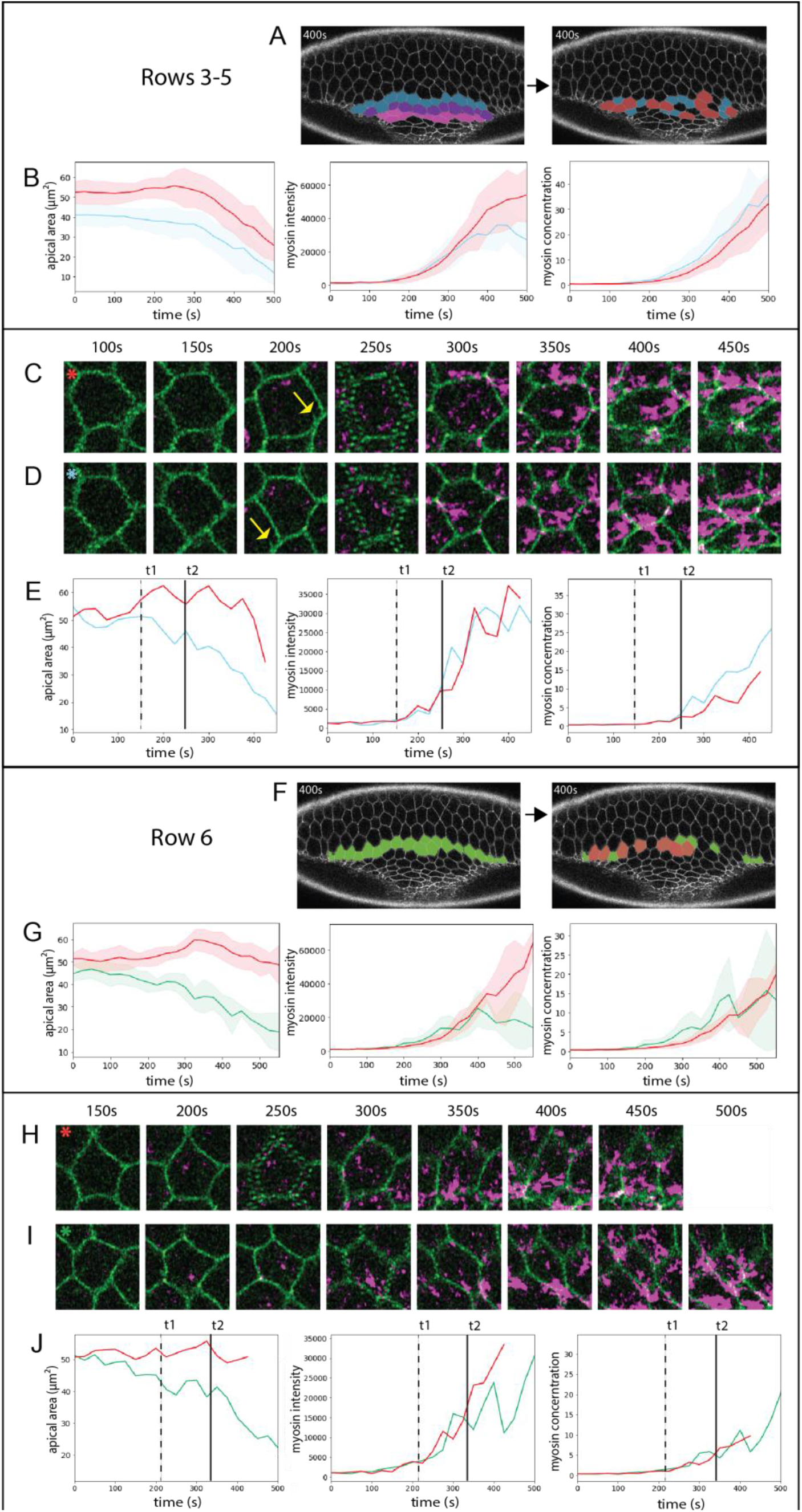
Myosin concentration in constricting and transiently expanding cells. Analysis of Embryo 1 expressing Spider::GFP (green) and Sqh::mCherry (magenta)(same as shown in Fig. 1E). Cells in the indicated rows were sorted into bins, defining cells as ‘transiently expanding’ (red) if they increased their apical areas by >10% of their initial area for at least 3 consecutive time points, and as contracting (blue for rows 3 to 5, green for row 6) if they decreased their apical areas over 10% of their initial area for at least 10 time points. (A – E) Constricting rows 3 – 5 (F – J) Transition row 6 (A, F) Image at t= 400s; left: colouring shows rows; right: colouring indicates the individual cells that were analysed (red, transiently expanding, blue or green, constricting). (B, G) Cell apical area, total myosin intensity and myosin concentration of constricting and transiently expanding cells plotted against time, shown as mean (solid line) and standard deviation (shaded area). (C - E) Analysis of a transiently expanding (C) and a constricting (D) cell from the constricting rows. (C, D) Snap shots of the two cells at the indicated time points. The two cells are adjacent to each other: the arrow at 200 sec points at a feature of the expanding cell that is also seen in the panel below. (E) Apical cell area, total myosin intensity and myosin concentration of the cells in C (red) and D (blue) plotted against time. t1 (dashed line) marks the divergence of the cells in apical area, t2 (solid line) the divergence in myosin concentration. (H - J) Analysis of a transiently expanding (H) and a constricting (I) cells. (H, I) Snap shots of the two cells at the indicated time points (J) Apical cell area, total myosin intensity and myosin concentration of the cells in H (red) and I (green) plotted against time. t1 (dashed line) marks the divergence of the cells in apical area, t2 (solid line) marks the divergence in myosin concentration.

## Supplementary videos

**Suppl. video 1.**
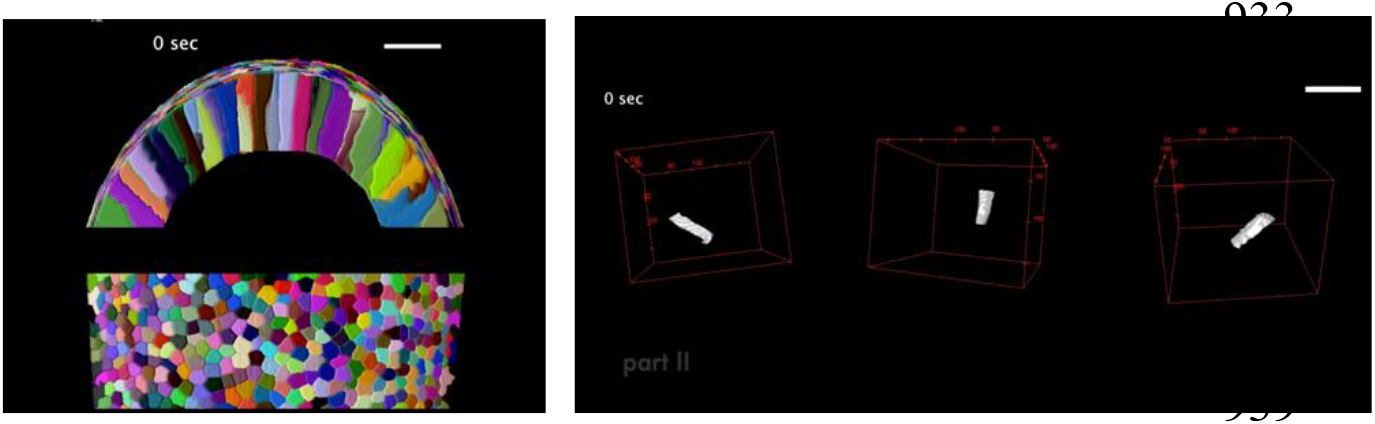
Part I. Cross sectional (top) and ventral (bottom) view of 3D segmented ventral half of an embryo expressing GAP43::mCherry and imaged with SPIM. Each colour marks a unique cell that is tracked in time. Part II. 3D volume rendering shown over time for 3 cells: one central and two (left and right) lateral mesodermal cells. The video illustrates the volume transited by the cells during ventral furrow formation. The tip of the left cell moved out of the imaging volume during the period. Apical is up. (scale bar = 20μm)

**Suppl. video 2.**
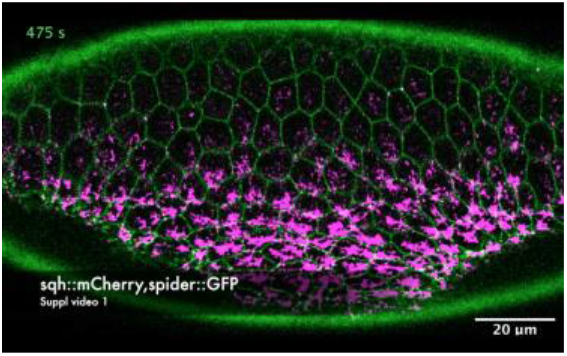
Ventro −lateral view of 2 embryos (parts I and II) expressing shq::mCherry (magenta; myosin) and Spider::GFP (green; membrane) showing the dynamics of apical area and myosin during ventral furrow formation and lateral cell expansion. t=0 in both movies is defined as 100 sec before first appearance of myosin in central mesodermal cells. 25 sec time steps. (Figs. 1 and 2)

**Suppl. video 3.**
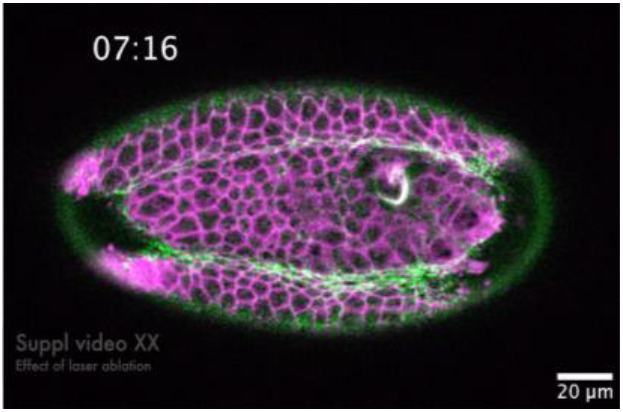
Ventral view of an embryo expressing GAP43mCherry (magenta; membrane), sqh::GFP (green; myosin). MCP::mCherry and Snail::MS2 (not shown) were used to mark the mesoderm boundary. The embryo is illuminated repeatedly in the area marked by red ellipse in frame 1. During the laser illumination experiment, images were captured every 2 sec. The last three frames are single confocal sections from a 3D stack taken at intervals of 38sec. (Fig. 3)

**Suppl. video 4.**
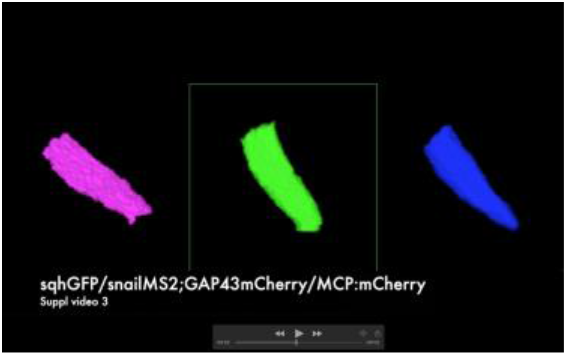
3D reconstructions of lateral mesodermal cells from row 7 in an embryo where apical constriction of the central mesodermal cells was inhibited by laser ablation. Apical is down, basal is up. Cells 1 and 2 fail to expand, cell 3 constricts apically. See main Fig. 3E-F for location of cells in the embryo.

**Suppl. video 5.**
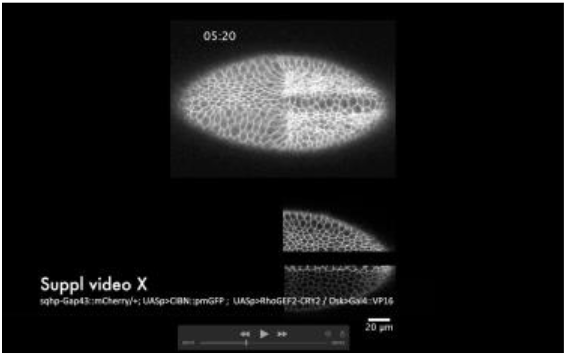
Ventral view of embryo expressing GAP43::mCherry (membrane, top), CIBN::pmGFP (membrane; bottom) and RhoGEF2-CRY2. Myosin is ectopically activated by illuminating the area in the yellow boxes in frame 1. (Fig. 4)

